# EXTRARNAS: A Framework for Extracting RNA Structures with Multiple Tools

**DOI:** 10.64898/2026.08.27.747497

**Authors:** Federico Di Petta, Piermichele Rosati, Piero Hierro Canchari, Michela Quadrini, Luca Tesei

**Affiliations:** School of Sciences and Technology, University of Camerino, Italy

**Keywords:** RNA structure annotation, base pairs, PDB, Docker, consensus structure

## Abstract

Accurate annotation of RNA base-pairing interactions is essential for structural analysis, benchmarking, and data-driven RNA structure prediction. Several tools can extract RNA interactions from three-dimensional coordinates, but their outputs are heterogeneous and may disagree, particularly for non-canonical base pairs. We present EXTRARNAS, a Java-based framework for automated, reproducible, and user-friendly large-scale extraction of RNA structural annotations with multiple tools. EXTRARNAS processes batches of RNA structures specified by PDB identifier and chain, or provided as local PDB files, executes annotation tools through a Docker-based environment, and parses tool-specific outputs using ANTLR4-based grammars. For each structure–tool pair, the framework generates standard BPSEQ files for canonical cis Watson–Crick interactions and introduces BPSEQE, a standardized text format for representing the extended secondary structure, preserving canonical, non-canonical, and multiple interactions per nucleotide. The current prototype supports RNAView, MC-Annotate, and RNAPolis Annotator. We demonstrate EXTRARNAS on eight RNA structures containing triple-helix motifs, comparing extracted canonical pairs against curated BPSEQ references and evaluating the recovery of manually validated Hoogsteen interactions. The results show consistent differences among tools, especially for non-canonical interactions, highlighting the need for standardized representations such as BPSEQE to support reproducible comparison and future consensus-based annotation.

## 1 Introduction

RNA molecules adopt complex structures whose biological functions depend on canonical and non-canonical interactions. Extracting base-pairing information from experimentally determined three-dimensional RNA structures is therefore a key step in structural bioinformatics, comparative analysis, and benchmarking of RNA structure-prediction methods. Several tools have been proposed for this task, including RNAView [1], MC-Annotate [2], RNAPolis Annotator [3], FR3D [4], X3DNA-DSSR [5], Barnaba [6], and BPNet [7].

A recent review on RNA structure prediction stresses that base-pair topology in public archives is often incomplete or unreliable and that different annotation programs may provide incomplete or conflicting information [8]. Moreover, most existing tools require independent installation, meaning that using multiple methods entails installing and maintaining each of them separately. Recent frameworks such as RNApdbee 3.0 [9] simplify this process by integrating multiple annotation tools and providing unified visualization of their results for individual RNA structures. However, constructing large, reproducible datasets for benchmarking and machine-learning applications still requires automated batch processing together with standardized textual representations of heterogeneous annotations. In particular, no standardized textual representation of the *extended secondary structure* is currently available across annotation tools, hindering systematic comparison and future consensus computation.

To address these limitations, we introduce **EXTRARNAS**, a Java-based framework for automated extraction and standardization of RNA structural annotations from three-dimensional data. EXTRARNAS provides a unified, reproducible, and user-friendly environment to execute multiple RNA annotation tools and convert their heterogeneous outputs into standardized representations suitable for direct comparison and downstream analysis. As summarized in Table 1, EXTRARNAS complements existing frameworks by focusing on standardized textual representations of RNA structural annotations. In particular, the framework generates standard BPSEQ files for canonical secondary structures and introduces BPSEQE, a standardized text representation of the *extended secondary structure* that preserves canonical, non-canonical, and multiple interactions in a tool-independent format.

**Table 1.** Comparison of the primary focus of EXTRARNAS and RNApdbee 3.0.

| Aspect | EXTRARNAS | RNApdbee 3.0 |
| --- | --- | --- |
| Typical execution | Automated batch processing | Interactive single-structure analysis |
| Primary goal | Annotation standardization and comparison | Integrated annotation and visualization |
| Representation of extended secondary structure | Single BPSEQE file | Multiple CSV reports |
| Consensus output | Planned structured consensus | Sequence-logo visualization |

The main contributions of this work are: (i) EXTRARNAS, a reproducible framework for batch execution, comparison, and standardization of RNA annotation tools, enabling systematic comparison of heterogeneous annotations and future consensus-generation strategies; and (ii) BPSEQE, a standardized text representation of the *extended secondary structure* that supports interoperable downstream analyses. We demonstrate the framework on RNA triple-helix structures, illustrating the systematic discrepancies that can arise across annotation tools.

## 2 Data and Methods

### Input data

EXTRARNAS takes as input a list of RNA structures specified by PDB identifier and chain identifier, provided as a CSV file. Each row of the file corresponds to a structure-chain pair. The framework automatically downloads the corresponding coordinate files from the Protein Data Bank and prepares tool-specific input files. If a local `.pdb` file with the same name as the input identifier is found in the shared working directory, EXTRARNAS uses the local file instead of downloading it. This design supports both large RNA datasets and custom or unpublished structures without manual retrieval or reformatting.

### Standardized output formats: BPSEQ and BPSEQE

Each annotation tool produces a different output format. EXTRARNAS uses ANTLR4-based parsers, generated from formal grammars of the supported formats, to extract base-pair annotations and convert them into a unified internal representation. For each input structure and annotation tool, EXTRARNAS generates two standardized outputs: BPSEQ, representing the RNA secondary structure, and BPSEQE, representing the *extended secondary structure*, including both canonical and non-canonical interactions.

The standard BPSEQ format consists of three space-separated columns (nucleotide index, nucleotide identity, and pairing partner) and represents only canonical cis Watson–Crick (cWW) base pairs. BPSEQE extends this representation by preserving the first two columns and replacing the pairing column with twelve columns corresponding to the considered Leontis–Westhof interaction classes:

~~~
id nt cWW tWW cWH tWH cWS tWS cHH tHH cHS tHS cSS tSS
~~~

Each interaction-class field contains either `0`, indicating that no interaction of that class exists for the nucleotide, or the index (or comma-separated indices) of the interacting nucleotide(s). By allowing multiple partner indices within the same interaction class, BPSEQE preserves all annotated interactions, including higher-order motifs such as base triples, without loss of information. Conflicting annotations produced by different tools are not merged. Instead, EXTRARNAS preserves the standardized BPSEQ/BPSEQE output generated by each annotation tool independently, enabling downstream comparison and future consensus extraction.

### Tool execution and reproducibility

RNA annotation tools may require different dependencies and execution environments. EXTRARNAS encapsulates each tool in a Docker-based execution layer (container), i.e., within lightweight containerized environments that bundle all required software dependencies. This allows tools to be executed consistently across different systems without manual installation.

Supported annotation tools are automatically deployed inside Docker containers, ensuring consistent execution across platforms without manual installation. The modular architecture simplifies the integration of additional annotation tools. The experiments reported in this paper were performed using EXTRARNAS v1.0.1-cibb2026, including the distributed Docker images, running on Java 21. The versions of the supported annotation tools are documented in the software repository (see Availability section).

### Architecture and implementation

Figure 1 illustrates the EXTRARNAS workflow and graphical interface. The system follows a modular architecture composed of two main components: a Java-based user interface developed with JavaFX and a Docker container that hosts the RNA annotation tools. Figure 1(a) summarizes the workflow. Starting from a CSV file containing PDB identifiers and chain IDs, EXTRARNAS retrieves RNA structures, executes the selected annotation tools within Docker, and converts their outputs into standardized BPSEQ and BPSEQE representations using ANTLR4-based parsers. The parsers were validated by successfully processing the complete outputs generated by the supported annotation tools for all RNA structures considered in this study and by manually verifying representative extracted annotations. Figure 1(b) presents the graphical user interface, which allows users to configure the workflow without interacting with command-line tools. Through the interface, users can load input datasets, select annotation tools, and specify extraction options. This design simplifies the use of multiple heterogeneous tools and makes the framework accessible to non-expert users.

**Figure 1.**
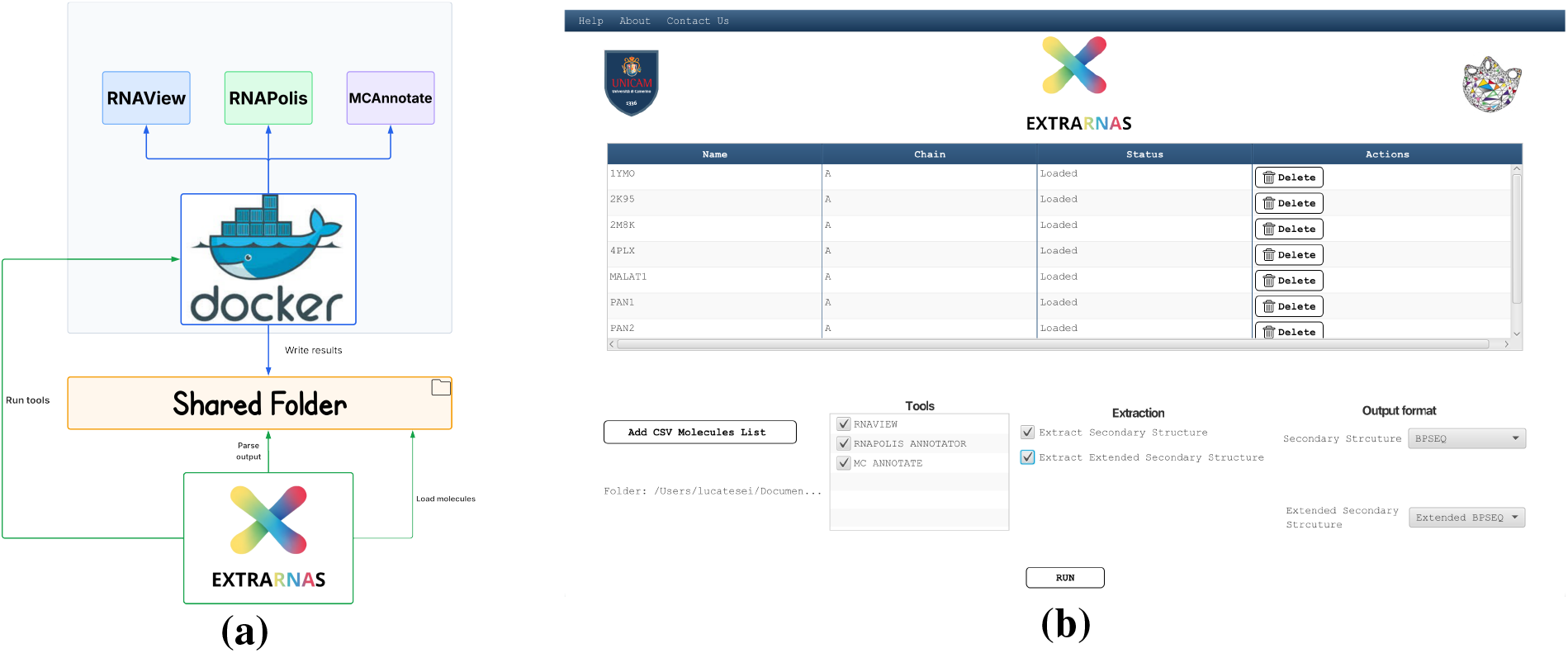
EXTRARNAS workflow and graphical interface. (a) Workflow of EXTRARNAS, from RNA structure retrieval to standardized BPSEQ and BPSEQE generation through Docker-based annotation tools. (b) JavaFX graphical interface for dataset loading, tool selection, and extraction workflow configuration.

Communication between the Java application and the Docker container is handled through the shared directory, which acts as a data exchange interface. The application automatically creates the Docker container when required, retrieves the supported annotation tools, executes them on the input dataset, stores the generated outputs in the shared working directory, and removes the container when processing is completed.

### Evaluation dataset and protocol

To provide a meaningful preliminary evaluation, we considered a small but manually curated dataset of RNA structures characterized by complex non-canonical interactions. In particular, we used a recently published validation dataset of RNA molecules containing triple-helix motifs and intricate Hoogsteen interactions [10]. Although intentionally limited in size, this dataset provides reliable reference annotations for both canonical base pairs and selected non-canonical interactions involved in triple-helix formation. Consequently, the reported results should be regarded as a preliminary demonstration rather than a comprehensive benchmark. For this short-paper evaluation, we selected the eight RNA molecules included in [10], namely `4PLX`, `2M8K`, `1YMO`, and `2K95` from the PDB, together with the `MALAT1`, `PAN1`, `PAN2`, and `hTER` structures provided by the authors as PDB files. For each structure, the dataset includes both a reference BPSEQ representation of canonical base pairs and a manually validated set of Hoogsteen interactions involved in triple-helix formation. These annotations allow us to evaluate the ability of RNAView, MC-Annotate, and RNAPolis Annotator to extract both canonical and non-canonical interactions.

Canonical base pairs are evaluated against the reference BPSEQ by computing true positives (TP), false positives (FP), false negatives (FN), and the corresponding F1-score. Non-canonical interactions are analyzed from BPSEQE files. In particular, Hoogsteen interactions involved in triple-helix motifs are evaluated only in terms of recovery of a manually validated subset. Since this reference includes only a curated subset of non-canonical interactions, pairs not present in the reference are not considered FP, and no precision or F1-score is computed for these interactions.

## 3 Results

### EXTRARNAS framework

EXTRARNAS provides an end-to-end workflow that generates standardized BPSEQ and BPSEQE outputs from batches of RNA structures processed by multiple annotation tools. The framework therefore provides reproducible execution of heterogeneous RNA annotation tools together with standardized BPSEQ and BPSEQE outputs for direct comparison. In particular, BPSEQE preserves non-canonical and multiple interactions in a unified representation across all supported annotation tools. Precompiled binaries, source code, Docker files, datasets, and analysis scripts are publicly available through the software and Zenodo repositories (see Availability section).

### Preliminary evaluation on triple-helix RNA structures

We applied EXTRARNAS to the eight RNA structures included in the curated triple-helix validation dataset of [10]. The goal of this preliminary evaluation was not to rank the tools, but to assess whether EXTRARNAS can make heterogeneous annotations directly comparable and expose systematic differences among annotators. Canonical and non-canonical interactions were evaluated as described in Section 2.

Table 2 reports aggregate results over the eight structures. The complete per-molecule comparison, including summary tables, tool-overlap tables, and missing/extra-pair lists, is provided in the associated Zenodo repository (see Availability section). F1 scores are computed per molecule and averaged across the dataset, while TP, FP, and FN are reported as totals. Non-canonical interactions are averaged per molecule, and triplex Hoogsteen interactions are reported as the number of recovered validated pairs over the total number of reference pairs.

**Table 2.** Aggregate evaluation on the eight triple-helix RNA structures. TP, FP, and FN are totals over canonical BPSEQ pairs, whereas precision, recall, and F1 are macro-averaged across molecules. NC avg. is the average number of non-canonical BPSEQE pairs per molecule. Triplex-H reports recovered manually validated triplex Hoogsteen pairs over the total reference pairs.

| Tool | Mol. | TP | FP | FN | Prec. | Rec. | F1 avg. | NC avg. | Triplex-H |
| --- | --- | --- | --- | --- | --- | --- | --- | --- | --- |
| MC_ANNOTATE | 8 | 145 | 4 | 29 | 0.974 | 0.852 | 0.894 | 6.250 | 31/51 |
| RNAPOLIS_ANNOTATOR | 8 | 152 | 8 | 22 | 0.951 | 0.887 | 0.916 | 6.875 | 33/51 |
| RNAVIEW | 8 | 126 | 5 | 48 | 0.941 | 0.780 | 0.821 | 6.875 | 21/51 |

The aggregate results highlight systematic differences among the annotation tools. RNAPolis Annotator achieves the highest average F1-score (0.916) and the lowest number of false negatives, indicating better coverage of canonical base pairs, although at the cost of a slightly higher number of false positives. MC-Annotate shows comparable performance (F1 = 0.894) with fewer false positives, suggesting a more conservative behavior. In contrast, RNAView exhibits a lower recall, as reflected by the higher number of false negatives (48), resulting in a lower F1-score (0.821).

Differences become more pronounced when considering non-canonical interactions. While all tools extract a comparable number of non-canonical pairs on average, their ability to recover manually validated triplex Hoogsteen interactions varies substantially. RNAPolis Annotator recovers 33 out of 51 validated pairs, followed by MC-Annotate (31/51), whereas RNAView recovers only 21/51. This confirms that the tools differ not only in the quantity but also in the type of non-canonical interactions they detect.

Overall, these results illustrate the type of discrepancies that EXTRARNAS is designed to expose. Even when canonical base-pair prediction is relatively consistent, substantial variability remains in the annotation of non-canonical interactions, particularly those involved in higherorder structural motifs such as triple helices. These discrepancies are partly due to differences in the geometric criteria used by each tool, as variations in distance and angular thresholds can lead to systematically different interaction annotations. This further supports the need for a framework that preserves tool-specific annotations while enabling systematic comparison and future consensus extraction.

## 4 Conclusion

We presented EXTRARNAS, a Java-based framework for automated, batch-oriented, and reproducible extraction of RNA structural annotations from three-dimensional data using multiple tools. EXTRARNAS integrates large-scale input processing, tool execution, parsing, and output normalization into a single workflow, producing standard BPSEQ files for canonical interactions and BPSEQE files for richer annotations including non-canonical and multiple interactions. Its Docker-based execution layer and JavaFX interface reduce installation complexity and make multi-tool RNA annotation accessible to non-expert users.

The preliminary evaluation on eight triple-helix RNA structures shows that EXTRARNAS can expose systematic discrepancies among annotation tools. While canonical base-pair extraction is relatively consistent, larger differences emerge for non-canonical interactions, in particular manually validated Hoogsteen pairs involved in triple-helix motifs. These results support the role of EXTRARNAS as a practical framework for comparative and large-scale RNA annotation, as well as a basis for future consensus extraction.

Future work will include the integration of FR3D, X3DNA-DSSR, Barnaba, and BPNet. Supporting an additional annotation tool requires implementing a dedicated ANTLR grammar together with a tool-specific execution wrapper, while the remaining workflow remains unchanged. Moreover, we will define automatic consensus strategies based on strict intersection, union, majority voting, weighted voting, and confidence-aware aggregation. Finally, we also plan to perform a systematic comparison of BPSEQE with existing textual formats for representing extended RNA secondary structures and a broader validation of its suitability for downstream RNA analysis workflows. These consensus structures may be evaluated against curated references and used to build higher-quality datasets for RNA structure prediction and benchmarking.

## Availability of data and software code

Software: https://github.com/bdslab/EXTRARNAS

Software release: https://doi.org/10.5281/zenodo.21238912

Datasets and scripts: https://doi.org/10.5281/zenodo.21246054

## Notes

https://doi.org/10.5281/zenodo.21246054

https://doi.org/10.5281/zenodo.21238912

https://github.com/bdslab/EXTRARNAS

### Competing Interest Statement

The authors have declared no competing interest.

### Funding

- European Union: National Recovery and Resilience Plan (NRRP), Mission 4, Component 2, Investment 1.1, under the PRIN 2022 PNRR call (Min. Decree No. 1409, dated September 14, 2022), project: P2022FFEWN RNA secondary structures and their relationship with function: application to non-coding RNAs (RNA2Fun), CUP: J53D23014960001

## References

[1] Huanwang Yang, Fabrice Jossinet, Neocles Leontis, L Chen, J Westbrook, H Berman, and Eric Westhof. Tools for the automatic identification and classification of RNA base pairs. Nucleic Acids Research, 31(13):3450–3460, 2003. DOI: 10.1093/nar/gkg680.

[2] Patrick Gendron, Sébastien Lemieux, and François Major. Quantitative analysis of nucleic acid three-dimensional structures. Journal of Molecular Biology, 308(5):919–936, 2001. DOI: 10.1006/jmbi.2001.4626.

[3] Marta Szachniuk. RNApolis: Computational platform for RNA structure analysis. Foundations of Computing and Decision Sciences, 44(2):241–257, 2019. DOI: 10.2478/fcds-2019-0012, Code: https://github.com/tzok/rnapolis-py.

[4] Michael Sarver, Craig L Zirbel, Jesse Stombaugh, Amani Mokdad, and Neocles B Leontis. FR3D: finding local and composite recurrent structural motifs in RNA 3d structures. Journal of Mathematical Biology, 56(1–2):215–252, 2008. DOI: 10.1007/s00285-007-0110-x.

[5] Xiang-Jun Lu, Harmen J Bussemaker, and Wilma K Olson. DSSR: an integrated software tool for dissecting the spatial structure of RNA. Nucleic Acids Research, 43(21):e142, 2015. DOI: 10.1093/nar/gkv716.

[6] Sandro Bottaro, Giovanni Bussi, Giacomo Pinamonti, Stefan Reisser, Wouter Boomsma, and Kresten Lindorff-Larsen. Barnaba: software for analysis of nucleic acid structures and trajectories. RNA, 25(2):219–231, 2019. DOI: 10.1261/rna.067058.118.

[7] Pritha Roy and Dhananjay Bhattacharyya. Contact networks in RNA: a structural bioinformatics study with a new tool. Journal of Computer-Aided Molecular Design, 36(2):131–140, 2022. DOI: 10.1007/s10822-021-00428-.

[8] Bohdan Schneider, Blake Alexander Sweeney, Alex Bateman, Jiri Cerny, Tomasz Zok, and Marta Szachniuk. When will RNA get its AlphaFold moment? Nucleic Acids Research, 51(18):9522–9532, 2023. DOI: 10.1093/nar/gkad726.

[9] J. Pielesiak, K. Niznik, P. Snioszek, G. Wachowski, M. Zurawski, M. Antczak, M. Szachniuk, and T. Zok. RNApdbee 3.0: A unified web server for comprehensive RNA secondary structure annotation from 3D coordinates. Journal of Molecular Biology, 438:169795, 2026. DOI: 10.1016/j.jmb.2026.169795.

[10] Margherita A G Matarrese, Michela Quadrini, Nicole Luchetti, Federico Di Petta, Daniele Durante, Monica Ballarino, Letizia Chiodo, and Luca Tesei. Decoding RNA triple helices: identification from sequence and secondary structure. Briefings in Bioinformatics, 27(1):bbag009, 2026. DOI: 10.1093/bib/bbag009.

